## Supplementary Information for "Cell-type-specific alternative splicing in the cerebral cortex and kidney of a *Setbp1^S858R^* Schinzel-Giedion Syndrome patient variant mouse"

| **Tissue** | **Cell type** | **Marker(s)** |
| --- | --- | --- |
| Cerebral Cortex | Neurons | *Snap25* |
| Cerebral Cortex | Excitatory Neurons | *Slc17a7* |
| Cerebral Cortex | Inhibitory Neurons | *Gad1, Gad2* |
| Cerebral Cortex | Microglia | *Cx3cr1, Cd68* |
| Cerebral Cortex | Astrocytes | *Slc1a3, Gja1* |
| Cerebral Cortex | Vascular Cells | *Vtn, Mgp, Bnc2, Pdgfrb, Col1a1, Dcn, Flt1* |
| Cerebral Cortex | OPCs | *Pdgfra* |
| Cerebral Cortex | Oligodendrocytes | *Mbp* |
| Kidney | B cells | *Cd79a. Cd79b* |
| Kidney | CDIC-A | *Aqp6* |
| Kidney | CDIC-B | *Hmx2* |
| Kidney | CDPC | *Aqp2, Hsd11b2* |
| Kidney | Connecting tubule cells | *Atp6v1b1, Fxyd4* |
| Kidney | DCT | *Slc12a3* |
| Kidney | Dendritic cells | *Itgax, H2-Aa* |
| Kidney | Endothelial cells | *Flt1* |
| Kidney | PCT | *Slc5a2, Slc6a19* |
| Kidney | PST | *Slc22a7, Slc22a6* |
| Kidney | Podocytes | *Nphs1, Synpo* |
| Kidney | Proximal tubule cells | *Slc34a1, Slc13a3* |
| Kidney | T cells | *Cxcr6, Cd247* |
| Kidney | T regulatory cells | *Ikzf2* |
| Kidney | Thick ascending limb (LOH) | *Ptger3, Tmem207* |
| Kidney | Thin ascending limb (LOH) | *Epha7, Mx2* |
| Kidney | Thin descending limb (LOH) | *Fst, Aqp1* |

**Table S1. Cell-type markers for cell-type assignment.** We used the above canonical marker gene expression to assign cell types.

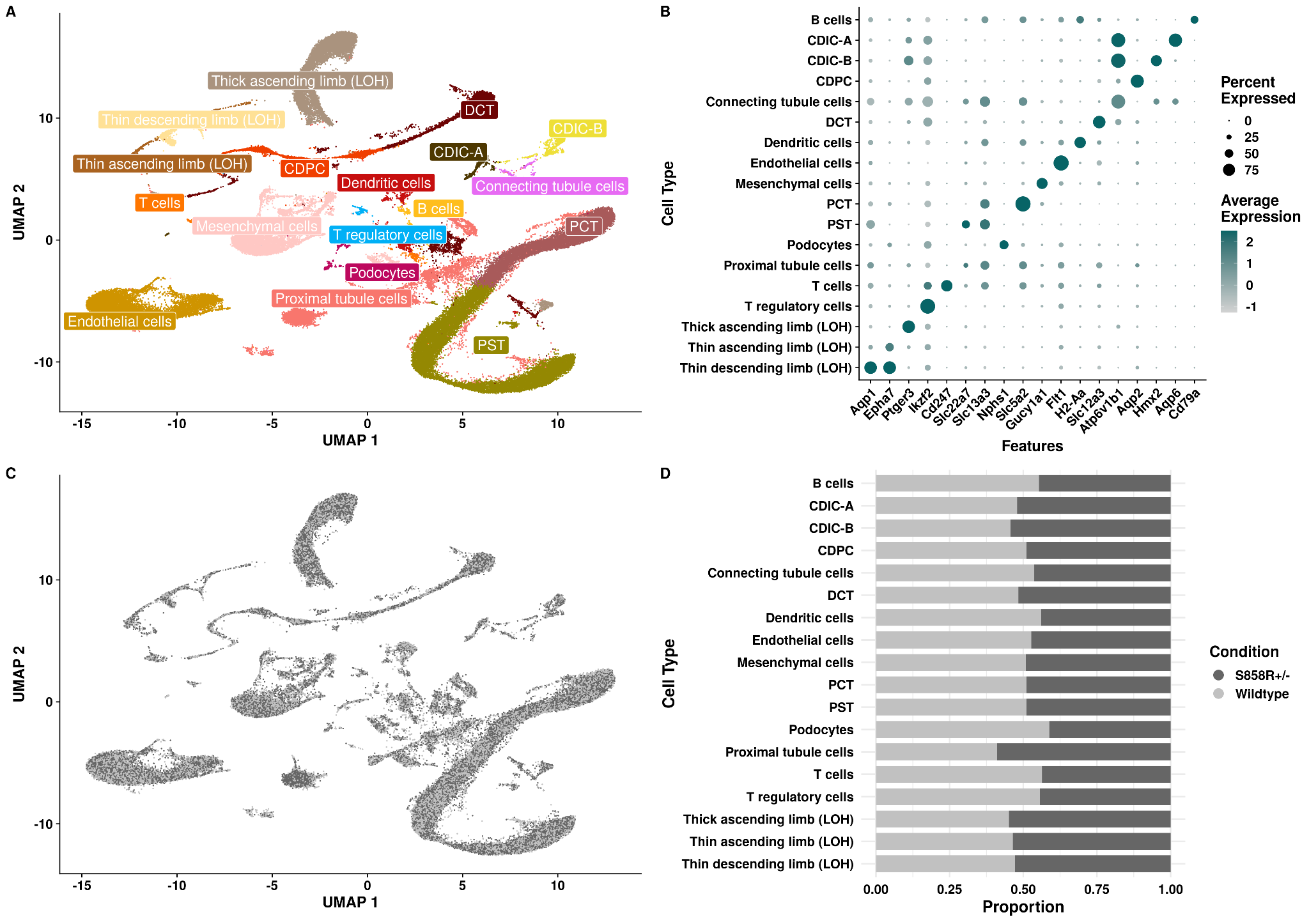

**Figure S1.** **snRNA-seq data overview of *Setbp1^S858R^* and wild-type kidney samples.** (A) Representative uniform manifold approximation and projection (UMAP) colored by cell types annotated in our integrated dataset. (B) A dot plot of representative cell type markers for cell types is shown in panel A. Size represents the percent of cells expressing a given gene, and color saturation represents average expression. (C) UMAP colored by condition. (D) Bar plot of cell type proportions split by condition. Dark gray denotes *Setbp1^S858R^* mice, and light gray denotes wild-type mice.

**Supporting Information Table 2. Number of SJs per gene per cell type in the cerebral cortex.** The table includes the number of SJs per gene per cell type for the 7 cerebral cortex cell types.

**Supporting Information Table 3. Number of SJs per gene per cell type in the kidney.** The table includes the number of SJs per gene per cell type for the 18 kidney cell types.

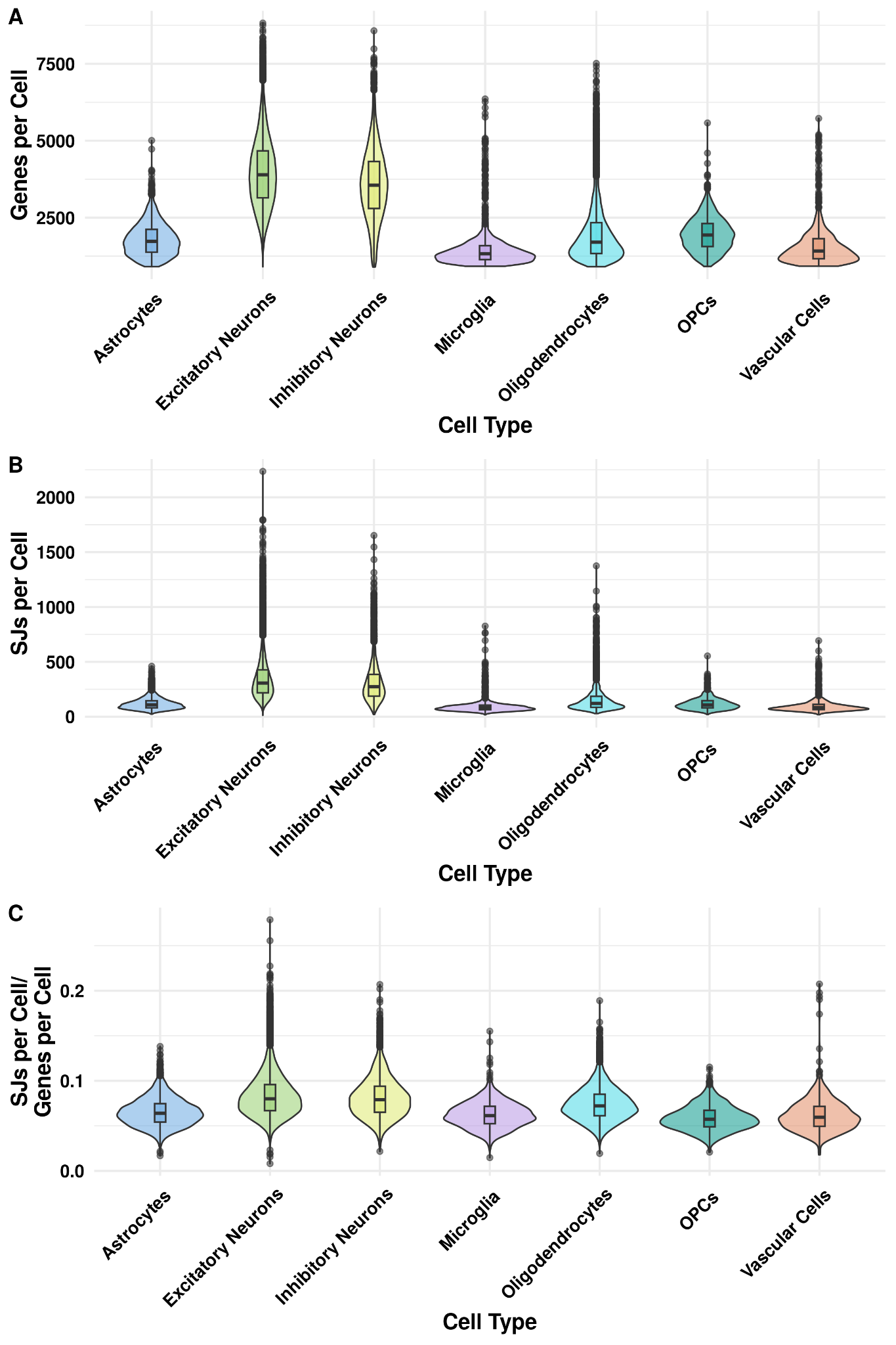

**Figure S2. Number of genes, splice junctions, splice junctions divided by genes expressed per cell split by cell type in the cerebral cortex.** Violin plots showing the number of (A) genes, (B) splice junctions, and (C) sj/genes expressed per cell split by cell type.

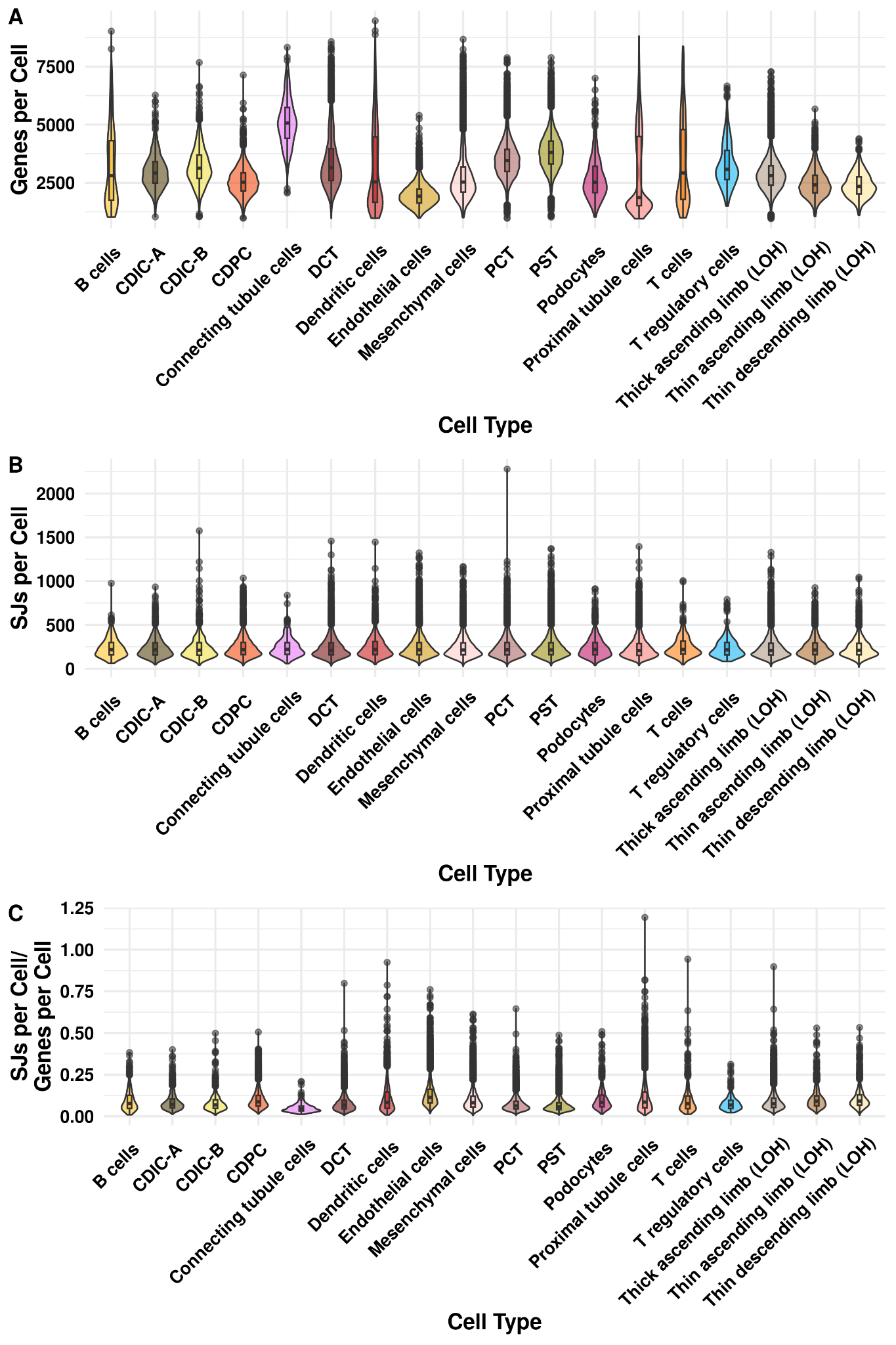

**Figure S3. Number of genes, splice junctions, splice junctions divided by genes expressed per cell split by cell type in the kidney.** Violin plots showing the number of (A) genes, (B) splice junctions, and (C) sj/genes expressed per cell split by cell type.

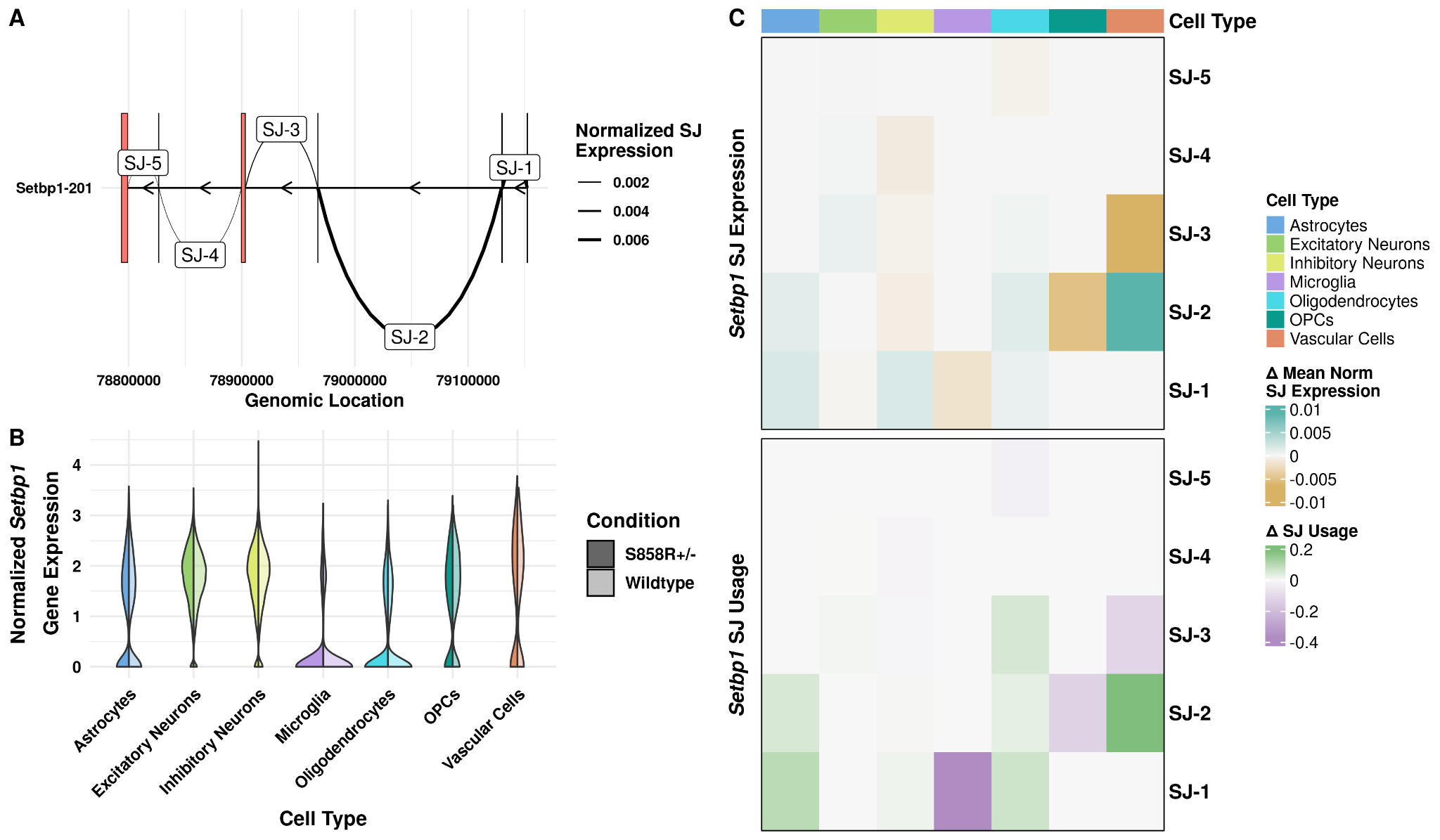
**Figure S4. *Setbp1* gene and SJ expression in the cerebral cortex.** (A) Transcript diagram showing the transcript structure of *Setbp1* in mouse. SJs are labeled on the curved lines connecting exons. Arrows indicate the direction of transcription (which for *Setbp1* is antisense). Connecting line thickness indicates the mean normalized SJ expression for a given SJ. (B) Split violin plots showing normalized *Setbp1* expression across cell type (x-axis) and condition (darker shade represents *Setbp1^S858R^* mice, and lighter shade represents wild-type mice). (C) Heatmaps showing the changes in normalized mean SJ expression (top) and usage (bottom) between *Setbp1^S858R^* and wild-type mice for the five different *Setbp1* SJs. The top annotation of columns shows cell type. A positive delta value indicates expression or usage was higher in the *Setbp1^S858R^* than in wild-type mouse brain tissue, and a negative indicates expression or usage was higher in wild-type mice.

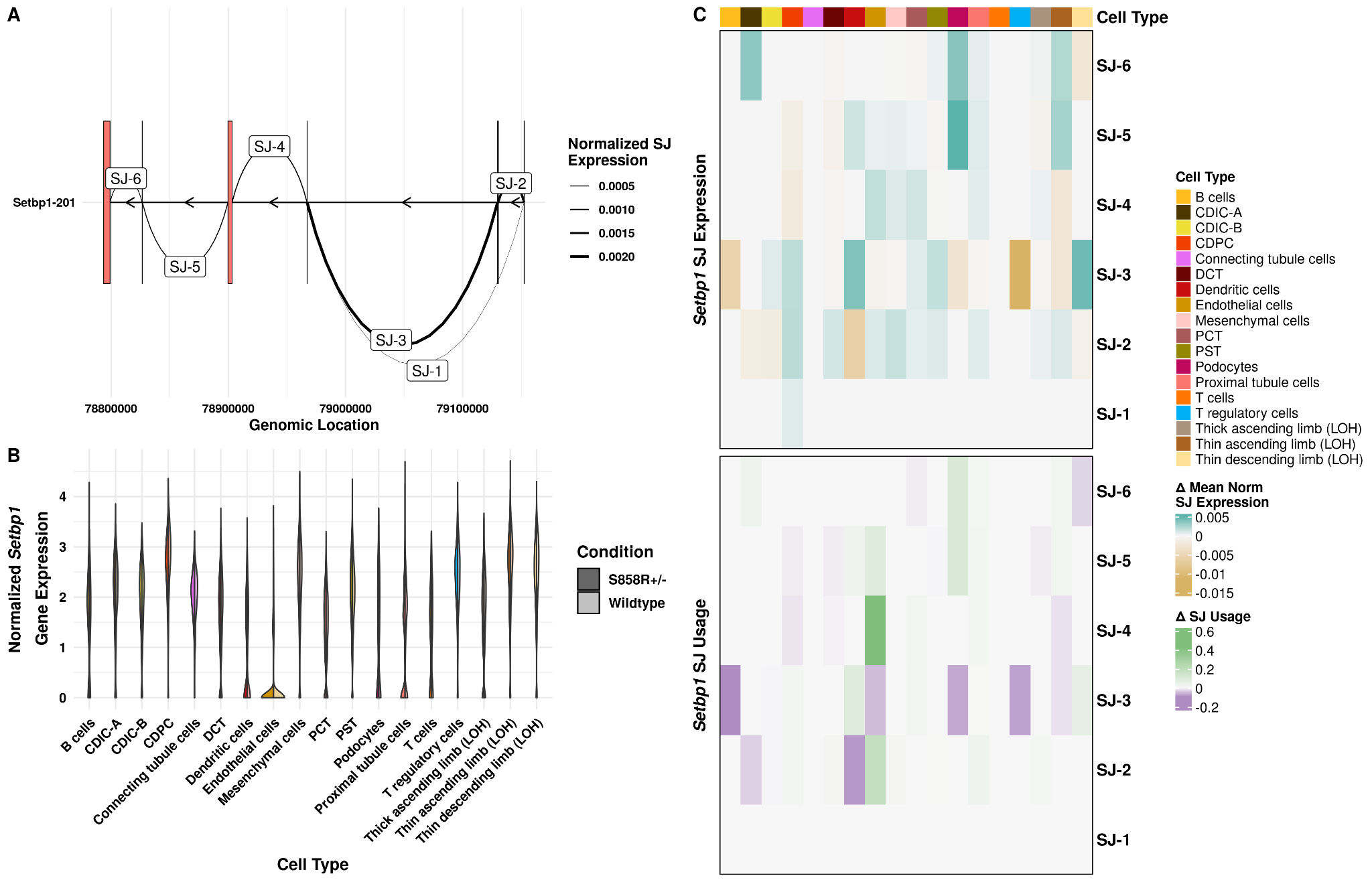

**Figure S5. *Setbp1* gene and SJ expression in the kidney.** (A) Transcript diagram showing the transcript structure of *Setbp1* in mouse. SJs are labeled on the curved lines connecting exons. Arrows indicate the direction of transcription (which for *Setbp1* is antisense). Connecting line thickness indicates the mean normalized SJ expression for a given SJ. (B) Split violin plots showing normalized *Setbp1* expression across cell type (x-axis) and condition (darker shade represents *Setbp1^S858R^* mice, and lighter shade represents wild-type mice). (C) Heatmaps showing the changes in normalized mean SJ expression (top) and usage (bottom) between *Setbp1^S858R^* and wild-type mice for the six different *Setbp1* SJs. The top annotation of columns shows cell type. A positive delta value indicates expression or usage was higher in the *Setbp1^S858R^* than in wild-type mouse brain tissue, and a negative indicates expression or usage was higher in wild-type mice.

**Table S4.** **Functional and disease annotations of genes with significant SJU.** Human-mapped ortholog gene symbols and Ensembl IDs of genes with significant SJUs (“Mouse”), annotated for GenCC disease titles (GenCC_Diseases), COSMIC CGD tier, COSMIC gene-associated somatic tumors, SFARI gene scores, SFARI genetic category, which tissue(s) (brain and kidney) we detected significant SJUs in, and the specific cell types in brain and kidney where we detected significant SJUs.

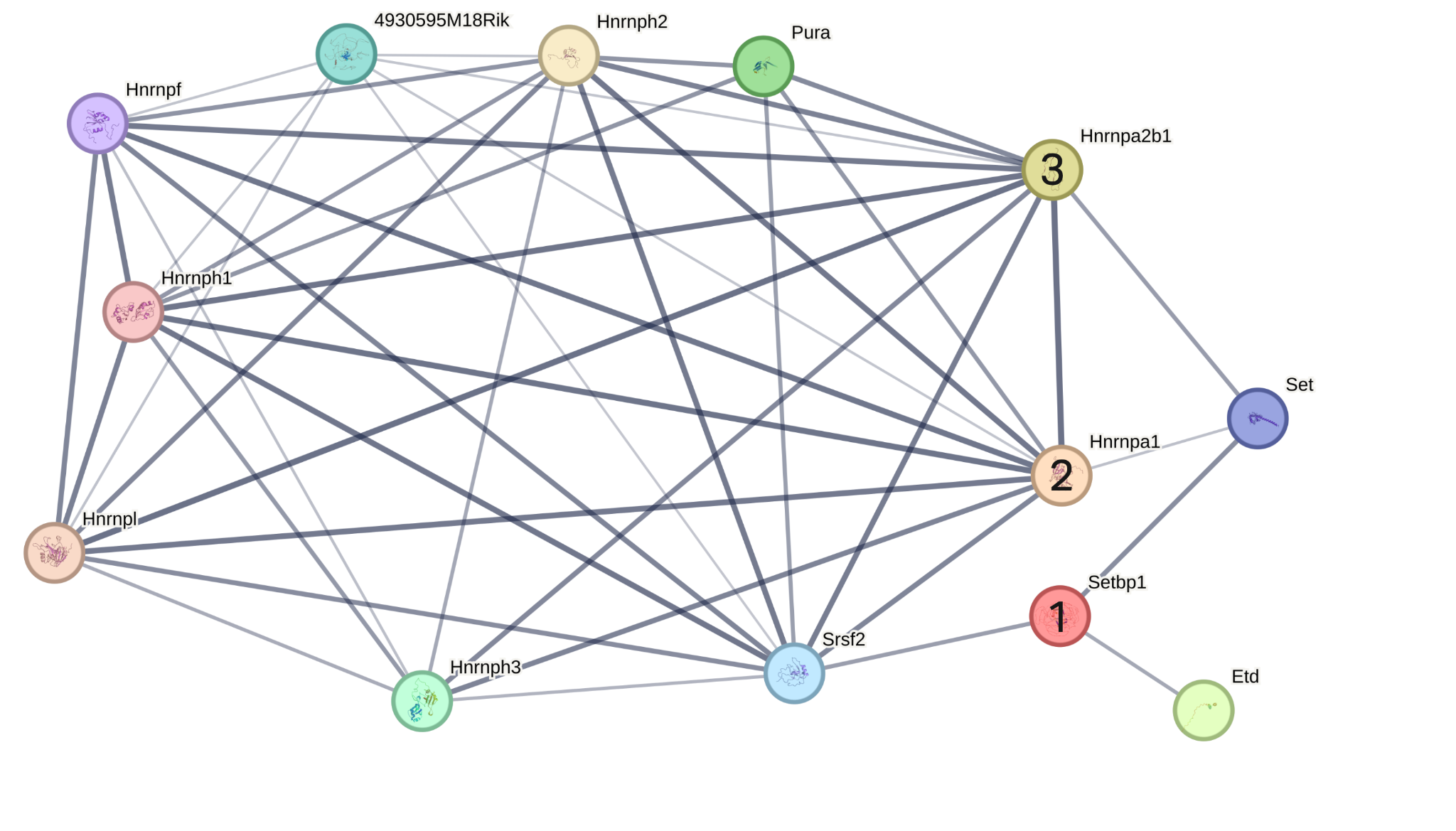

**Figure S6. StringDB protein-protein interaction (PPI) network.** PPI of Setbp1 (node 1), Hnrnpa1 (node 2), and Hnrnpa2b1 (node 3). Edges represent interactions with known evidence compiled in StringDB. Thicker edges represent higher confidence scores.

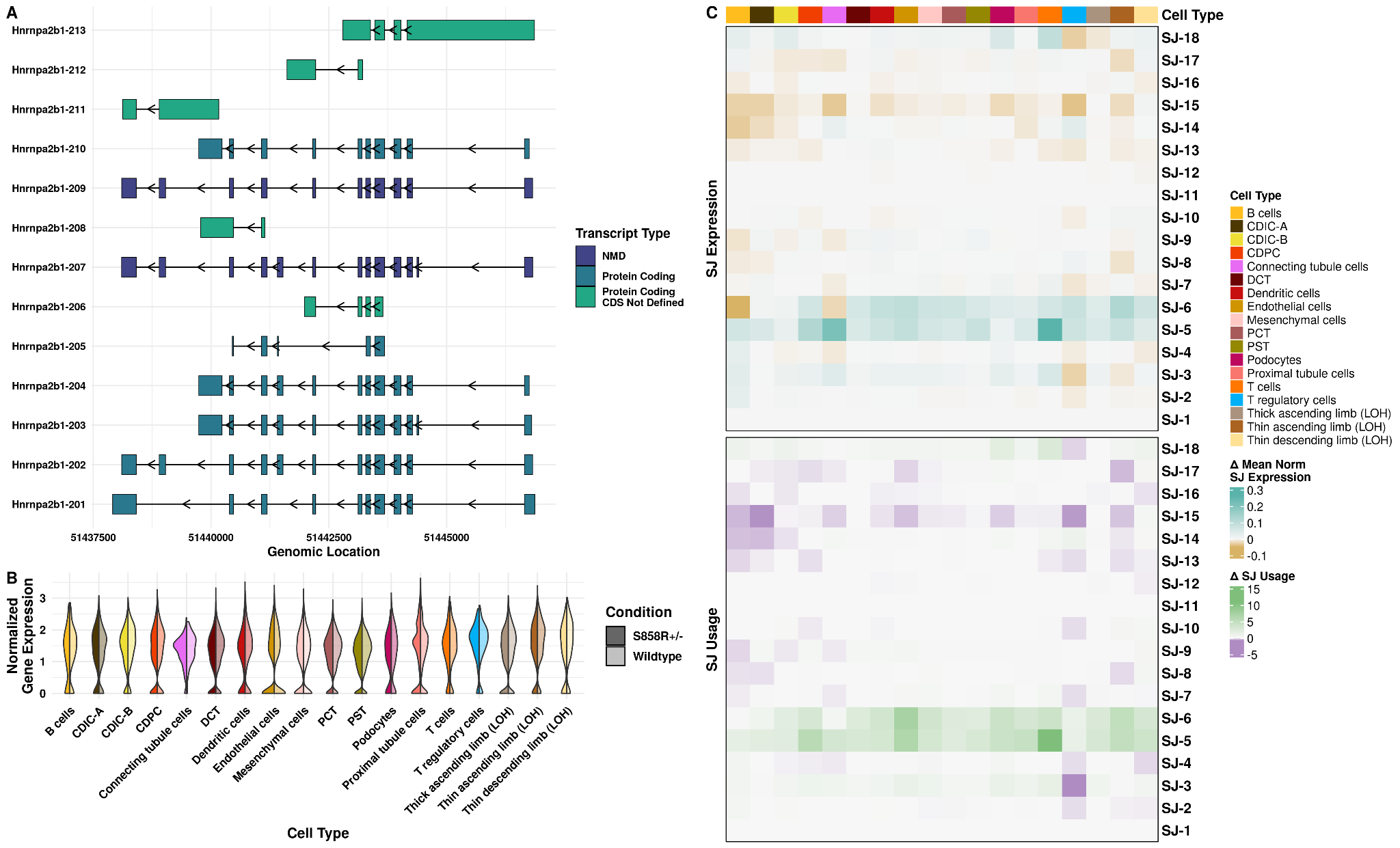

**Figure S7. *Hnrnpa2b1* has significant AS changes in 12 of 18 kidney cell types in *Setbp1*^S858R^ compared to wild-type mice.** (A) Transcript models of all 13 annotated transcripts of *Hnrnpa2b1*. The color indicates transcript classification: indigo = transcripts flagged for nonsense-mediated decay (NMD), dark teal = protein-coding transcripts, turquoise = protein-coding transcripts, but coding sequence (CDS) is not defined. Arrows indicate the direction of transcription. (B) Split violin plots showing *Hnrnpa2b1* normalized gene expression per cell for all cell types, split by condition. (C) Heatmaps of the changes in normalized mean SJ expression (top) and usage (bottom) between *Setbp1^S858R^* and wild-type mice for 18 SJs of *Hnrnpa2b1*. The top heatmap annotation indicates cell type. A positive delta indicates expression or usage was higher in *Setbp1^S858R^* than in wild-type mice, and a negative indicates expression or usage was higher in wild-type mice.
